# PA-SfM: Tracker-free freehand 3D photoacoustic imaging via differentiable acoustic radiation

**DOI:** 10.64898/2026.04.06.716718

**Authors:** Shuang Li, Jian Gao, Chulhong Kim, Seongwook Choi, Hao Huang, Xuanhao Wang, Junhui Shi, Qian Chen, Yibing Wang, Shuang Wu, Yu Zhang, Yucheng Zhou, Yao Yao, Changhui Li

## Abstract

Three-dimensional photoacoustic imaging (3D PAI) has shown great potential in preclinical studies and clinical implementations, while it commonly has sparse sensor arrays, limiting both spatial sampling and the field of view (FOV). Freehand scanning could overcome these limitations through anatomy-adaptive scanning that extends spatial coverage and provides complementary views beyond the fixed array aperture, but its flexibility is conventionally constrained by the need for motor feedback or external pose tracking. Here we introduce PA-SfM, a tracker-free structure-from-motion (SfM) framework that recovers multiple poses directly from PA measurements with superior precision, enabling freehand 3D PAI without external pose measurements. PA-SfM integrates a differentiable acoustic radiation model with hierarchical optimization and rigid array constraints, it jointly estimates inter-view transformations and reconstructs 3D volumes. The performance of PA-SfM was demonstrated in numerical simulations, animal studies and human imaging experiments. Compared with motor encoder-based fusion of the mechanically scanned system, PA-SfM produced sharper and more continuous vascular reconstructions with an expanded FOV. In controlled quantitative validations, PA-SfM achieved high reconstruction fidelity, with PSNR values over 40 dB and SSIM values above 0.98 relative to ground truth or known pose reference reconstructions. These results demonstrate that PA-SfM enables robust freehand 3D PAI, with particular potential for clinical implementation of large-scale freehand imaging. The source code is publicly available at https://github.com/JaegerCQ/PA-SfM.

## 1 Introduction

Photoacoustic imaging (PAI) is a hybrid imaging modality that combines optical absorption contrast with ultrasound detection, and is widely explored in both pre-clinical and clinical settings [1–6]. Technological advances have greatly facilitated three-dimensional (3D) PAI based on two-dimensional matrix arrays, including spherical and planar arrays [7–14], as well as synthetically scanned systems [15, 16]. Owing to these advantages, 3D PAI has successfully demonstrated its superior *in vivo* imaging of peripheral vessels [15, 17], breast tissue [18, 19], and small animals [20].

Despite these advances, sensor arrays in 3D PAI are often sparse because of physical and engineering constraints. As a result, a single acquisition pose may provide insufficient spatial sampling density or angular coverage for high quality volumetric reconstruction [21, 22]. Acquiring data from multiple views is therefore a common strategy to increase both the sensor density and effective acquisition aperture, mitigating reconstruction artifacts. In practice, these additional views are often generated through mechanical scanning consisting of rotation and translation [1, 15, 23, 24]. The success of multi-view reconstruction critically depends on accurate knowledge of the sensor pose across views. Even minor positioning errors can lead to structural blurring, vessel discontinuity, and loss of quantitative fidelity in the reconstructed volume.

Various external tracking mechanisms are employed to provide physical pose feed-back. In controlled systems, motorized stages and mechanical encoders are used to trace the acquisition trajectory. In freehand settings, external positioning devices such as optical tracking systems, electromagnetic trackers, and inertial measurement units are commonly employed [25, 26]. Although these methods can provide geometric information, they also increase system complexity, cost, and calibration burden. Optical tracking requires a clear line of sight that is often difficult to maintain in crowded experimental or clinical environments [25, 27]. Electromagnetic tracking is susceptible to field distortion from metallic objects and nearby electronics [26]. Inertial sensing is portable and inexpensive, but its estimates are vulnerable to drift accumulation over time [25, 28]. Mechanical scanning systems offer controlled motion, yet the final reconstruction remains limited by encoder precision, assembly tolerance, and motion induced vibration.

These limitations have motivated increasing interest in tracker-free or sensorless pose estimation [29–31]. However, this problem is fundamentally challenging in photoacoustic imaging. Unlike ultrasound, photoacoustic signals do not exhibit stable speckle patterns that support standard speckle tracking or cross-correlation based motion estimation [32–35]. Deep learning approaches can learn motion priors from reconstructed images, but they require large labeled datasets and may not generalize across imaging systems or acquisition conditions [36]. Other image-based strategies, including frequency domain analysis of vascular patterns [37], can estimate relative probe motion only under restricted conditions. In particular, they typically require the displacement between adjacent frames to remain within 0.5 mm to 2 mm and the overlap between views to exceed 85%, which greatly limits the practicality of freehand scanning when displacement is large, structural features are weak, or probe motion changes abruptly. To address these challenges, we present PA-SfM, a tracker-free framework for free-hand 3D PAI with joint pose recovery and reconstruction in multi-view, expanded field of view (FOV). PA-SfM estimates relative imaging poses directly from PA signals by combining a differentiable acoustic radiation model with hierarchical optimization under rigid-array geometry constraints. Starting from the initial view, PA-SfM localizes the sensor array in subsequent views, enforces rigid-array consistency to obtain coarse relative poses, and further refines the global transformations through differentiable pose optimization before joint multi-view or multi-pose reconstruction. We demonstrate genuine freehand 3D PAI of human hand vasculature, where arbitrary hand motion provides multi-view measurements for pose recovery and joint reconstruction of a large FOV vascular network based on PA-SfM without external motion tracking. In this freehand experiment, the maximum average probe displacement between adjacent poses exceeded 9 mm, highlighting the strong practicality and robustness of PA-SfM under large and uncontrolled probe motion. We further validate the method in numerical simulations, *in vivo* rat experiments with known relative geometry, and a mechanically scanned 3D PAI system operated with both rotation and translation. In mechanically rotated imaging, PA-SfM reduced inter-view misregistration and produced sharper, more continuous vascular reconstructions than joint reconstruction based on motor encoder. In translational multi-pose imaging, PA-SfM enabled expanded FOV PA mapping without using translation stage pose feedback. Together, these results establish PA-SfM as a general computational framework for tracker-free pose recovery and joint reconstruction in multi-view, expanded FOV and freehand 3D PAI.

## Results

We first demonstrated genuine freehand 3D PAI of human hand vasculature, where we freely moved the hand in a 3D PAI system to enable multi-view imaging with expanded FOV. We next compared the performance of PA-SfM and hardware tracking in mechanically rotated PAI system with position encoding. Then, we demonstrated effective expansion of the PAI system’s FOV via translational multi-pose acquisition without external pose feedback. Finally, we performed controlled validation in both numerical simulations and *in vivo* rat experiments with known relative geometry to quantify sensor localization, pose estimation and reconstruction accuracy.

### 2.1 PA-SfM freehand 3D reconstructions of hand vessels

To evaluate the feasibility of PA-SfM, we performed tracker-free freehand PAI of a 26- year-old male volunteer’s hand using the 3D-PanoPACT system [23]. 3D-PanoPACT is a hemispherical array PAI system with 1024 transducer elements and a diameter of 20 cm, providing volumetric acoustic detection over a large angular aperture. The central frequency of each transducer element is 3.16 MHz. The human study has passed the ethical approval of the Peking University Institutional Review Board. More details regarding the system’s configuration and imaging performance can be found in the literature [23].

In this freehand experiment, the hand was placed above the imaging aperture and freely moved without external tracking, while the PAI system is fixed, as shown in Figure 1(a). The hand motion included extension, retraction and swinging trajectories, thereby producing a sequence of partially overlapping photoacoustic views with unknown relative poses. PA-SfM was then used to estimate the inter-view transformations directly from the photoacoustic signals and to recover the absolute coordinates for each individual view, thus integrating into an equivalent large array for joint reconstruction. The results were reconstructed using the classical delay-and-sum (DAS) algorithm, and maximum-amplitude projections (MAPs) were used for visualization. Figure 1 illustrates the overall workflow. Compared with conventional static imaging, freehand acquisition provides multiple observations of the target from different relative viewpoints, but also introduces unknown hand motion and view-dependent misalignment. Fig. 1(b) presents the imaging geometry from top and side views, and Fig. 1(c) is a photography of experimental configuration. The freehand hand imaging experiment procedure can be found in Supplementary Movie 1. Although the volumetric frame rate is 10 Hz, to demonstrate the capability of PA-SfM to recover pose without much overlap in FOV, five single views were selected from over ten candidate views in chronological order as shown in Fig. 1(d), and each individual view captured only partial vascular information and exhibited view-dependent incompleteness. After PA-SfM pose recovery and joint reconstruction, the reconstructed volume reveals a more complete 3D vascular network of the hand with a substantially expanded FOV of 50 mm *×*30 mm *×*10 mm and fewer artifacts, as demonstrated by the top-, side- and front-view MAPs and the volumetric display in Fig. 1(e). The 3D display of the result can also be found in Supplementary Movie 2. Notably, the probe displacement between adjacent poses can exceed 9 mm, yet the PA-SfM reconstruction still preserved fine vascular details without visible vessel duplication or discontinuity. These results demonstrate that PA-SfM successfully recovered coherent 3D vascular morphology from freehand PA measurements without relying on external tracking methods.

**Fig. 1.**
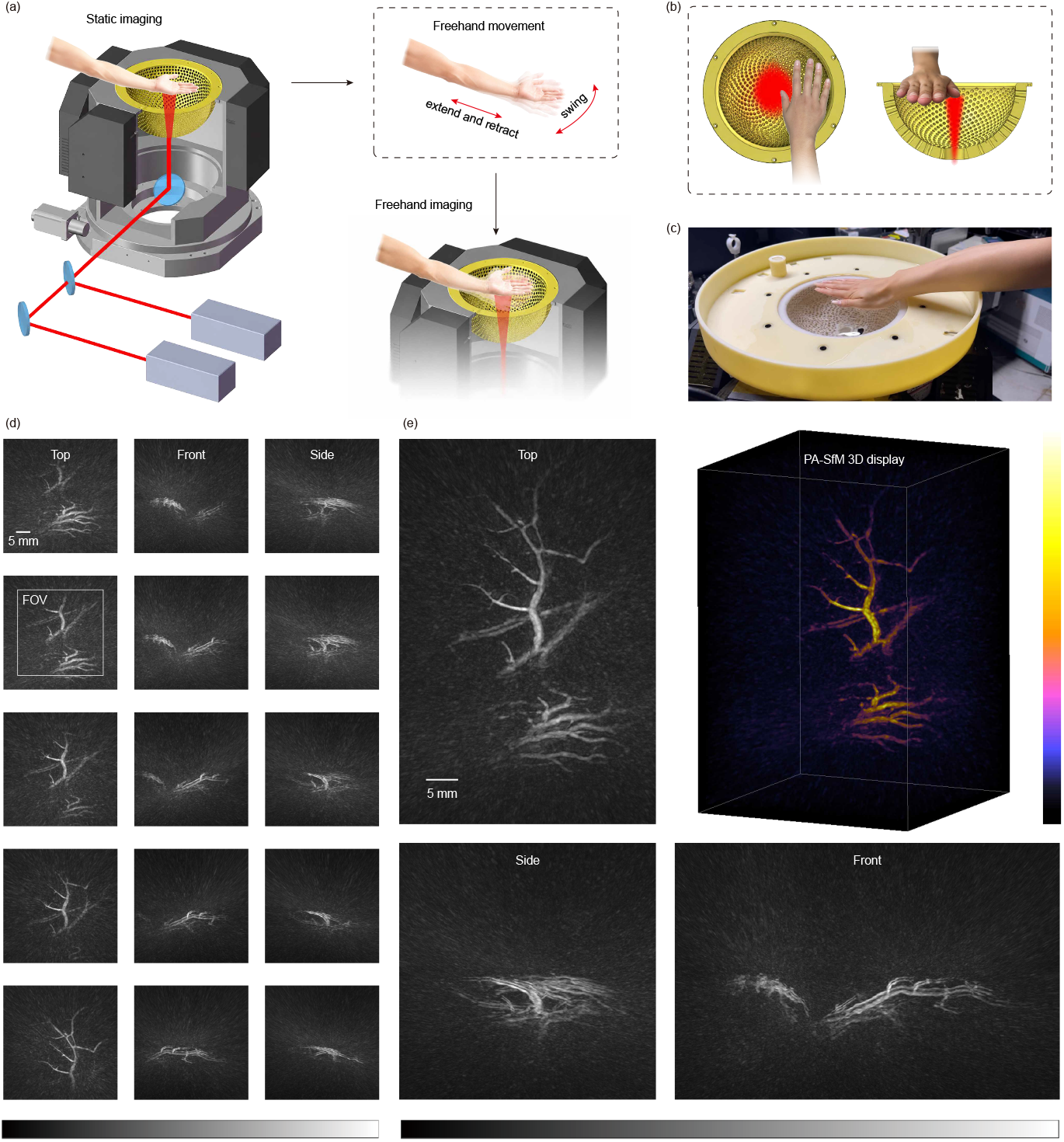
PA-SfM freehand 3D reconstructions of human hand. (a) Workflow of freehand photoacoustic imaging of the hand using the 3D-PanoPACT system. During imaging, the hand performed freehand motion, including extension–retraction and swinging trajectories, to provide multi-view measurements for PA-SfM reconstruction. (b) Detailed top and front views of the imaging geometry, illustrating the hand position, detection aperture and optical excitation region. (c) Photograph of the experimental setup. (d) Top-, front-, and side-view maximum amplitude projections (MAP) of reconstructions from five sequentially ordered single views (Scale: 5 mm; grayscale; reconstruction with DAS), in which the white box represent the FOV of a single view. (e) Top-, front-, side-view MAP and 3D display for PA-SfM freehand 3D reconstruction result (Scale: 5 mm; colormap: gray, magma).

To assess the repeatability of PA-SfM under freehand acquisition, we performed two additional independent hand imaging experiments, as shown in Fig. 2. In the first experiment, eight sequentially ordered but non-consecutive frames (fewer overlaps between frames) from imaging area 1 were selected for PA-SfM reconstruction. These frames were from the same freehand data used in Fig. 1, with three additional frames included to increase the spatial coverage and provide more diverse viewing angles. As shown in Fig. 2(a), PA-SfM successfully recovered a coherent vascular structure from these sparsely sampled views.

**Fig. 2.**
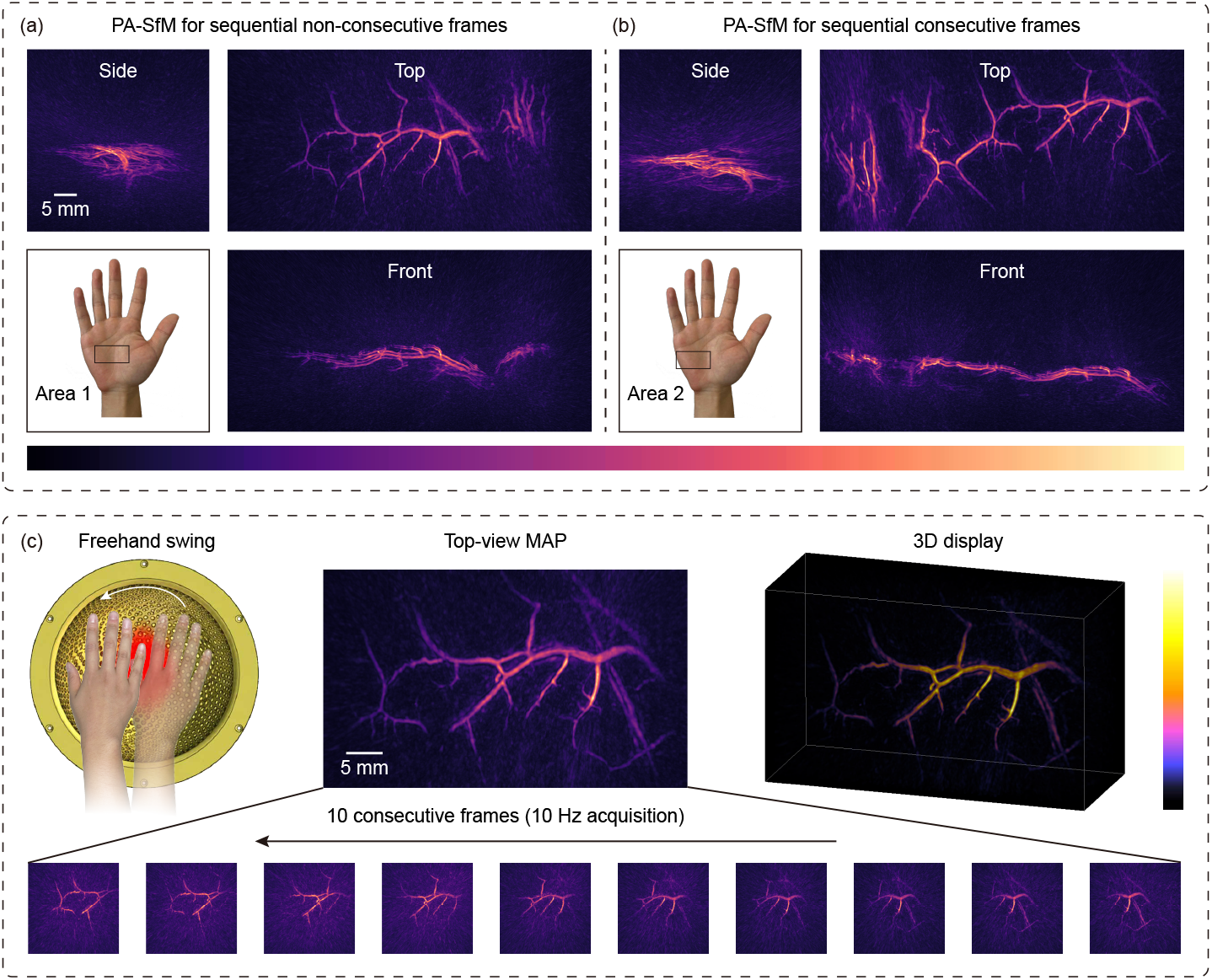
Repeatability validation of PA-SfM freehand 3D reconstruction of hand vessels using two partially overlapping imaging regions. (a) PA-SfM reconstruction of imaging area 1 using eight sequentially ordered but non-consecutive frames acquired from the same freehand hand vessel data as in Fig. 1, with three additional frames included (Scale: 5 mm; colormap: magma). (b) PA-SfM reconstruction of another hand vessel data acquired from imaging area 2 using 25 consecutive frames during a freehand swing trajectory ((a) and (b) are independent experiments derived from the same subject). (c) Pose recovery and joint 3D reconstruction visualization using the first 10 consecutive frames from (b) (Scale: 5 mm; colormap: magma).

In the second experiment, an independent freehand acquisition was performed on a different imaging area 2 of the same palm. This data consisted of 25 consecutive frames acquired (10 Hz sampling rate) during a freehand swing trajectory, providing a denser multi-view acquisition of the hand vasculature. PA-SfM reconstruction from these consecutive frames yielded a continuous 3D vascular map with consistent vessel morphology across the reconstructed FOV, as shown in Fig. 2(b). Compared with Fig. 2(a) and Fig. 2(b), there is no noticeable image quality difference, indicating that the method can tolerate moderate temporal discontinuity when sufficient structural overlap is preserved.

Since two independent imaging experiments have a partially overlapping area, we can validate the repeatability by comparing the reconstruction results in the overlapped area. Fig. 2(c) shows the PA-SfM pose recovery and joint reconstruction using the first 10 consecutive frames from the second experiment, which corresponds to the region overlapping with area 1. The major vascular branches and local vessel patterns reconstructed from the two experiments show consistent morphology in the shared region, despite being acquired from different frame selections and motion trajectories. The structural consistency observed in the partially overlapping regions further supports the robustness of the estimated poses and the reliability of the joint reconstruction. These findings indicate that PA-SfM can accommodate practical freehand imaging conditions, where frame overlap, motion continuity and viewing geometry may vary.

### 2.2 PA-SfM validation in animal study with rotational multi-view reconstruction

We next evaluated the performance of PA-SfM using *in vivo* animal experimental data [23] acquired with the same 3D PAI system for hand imaging in section 2.1. As aforementioned, it is a routine way to mechanically rotate the transducer array to increase the effective spatial sampling density and reduce limited-view artifacts. However, the final reconstruction quality is highly sensitive to the accuracy of mechanical positioning feedback. Hardware-induced errors, including encoder inaccuracies, mechanical structural deviations and stochastic vibrations during motor movement, can introduce inter-view misalignment and thereby degrade multi-view reconstruction.

The *in vivo* animal study is illustrated in Fig. 3(a), where the mouse hepatic vasculature was imaged from two views controlled by a mechanical 8.0° rotation of the hemispherical array around the system’s central *Z* axis. To compare PA-SfM with encoder-based tracking, multi-view joint reconstructions using either hardware feedback or PA-SfM are presented in Fig. 3.

**Fig. 3.**
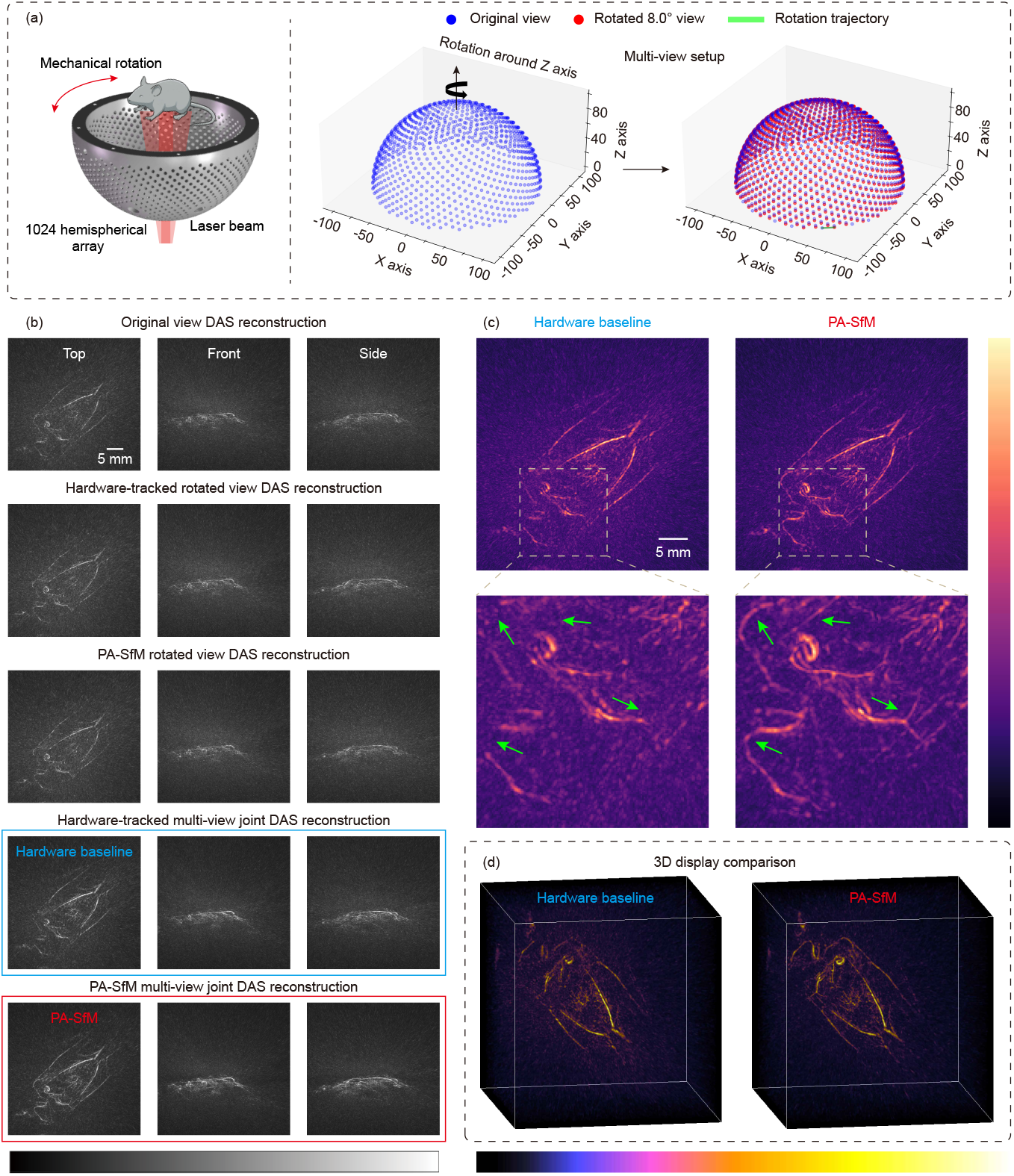
Multi-view PA reconstructions of mouse hepatic vasculature. (a) 3D PAI imaging system based on a mechanically rotating hemispherical array. (b) Top-, front-, and side-view MAPs of reconstructions from single views (original, hardware-tracked and PA-SfM rotated) and multi-views (hardware-tracked baseline, and PA-SfM result) (Scale: 5 mm; grayscale; reconstruction with DAS). (c) Locally magnified top-view MAPs comparing the hardware-tracked baseline and PA-SfM fine-tuned reconstruction (Scale: 5 mm; colormap: magma). (d) 3D display comparing hardware-tracked and PA-SfM reconstructions.

As shown in Fig. 3(b), both the original and rotated single-view reconstructions provide the same FOV with different fine vascular structures due to sparse spatial sampling. Without external hardware tracking, the PA-SfM reconstruction estimates the relative transformation directly from the photoacoustic data and successfully yields vasculature with rich fine details. It is worth noting that PA-SfM reconstruction presents markedly sharper and more continuous vascular structures than the reconstruction result based on the hardware-tracked mechanism. In the zoomed-in MAPs in Fig. 3(c), the green arrows highlight misregistered and fragmented vessels in the hardware-tracked reconstruction, whereas the PA-SfM reconstruction resolves these local discontinuities and recovers the microvascular morphology with much better vessel continuity. The 3D comparison in Fig. 3(d) further confirms that PA-SfM can achieve high quality multi-view reconstruction without hardware tracking.

Again, to validate the repeatability of PA-SfM, we chose another set of data that imaged a different part of the mouse abdomen with a nominal rotation angle of 4.0° around the system’s *Z* axis. As shown in Fig. 4, PA-SfM again improves the spatial consistency of the multi-view reconstruction compared with the hardware-tracked baseline. Fig. 4(a) presents two single-view reconstructions and the joint reconstruction result. From the zoomed-in images in Fig. 4(b), compared with hardware-tracked way, the PA-SfM reconstructed much finer vascular structures with prominent better continuity and less blurring, as marked by green arrows. These two comparisons with mechanical rotation motion indicate that PA-SfM achieves higher positioning accuracy than this system’s hardware tracking, which could be influenced by multi-ple factors, such as mechanical structural deviations and stochastic vibrations during motor movement.

**Fig. 4.**
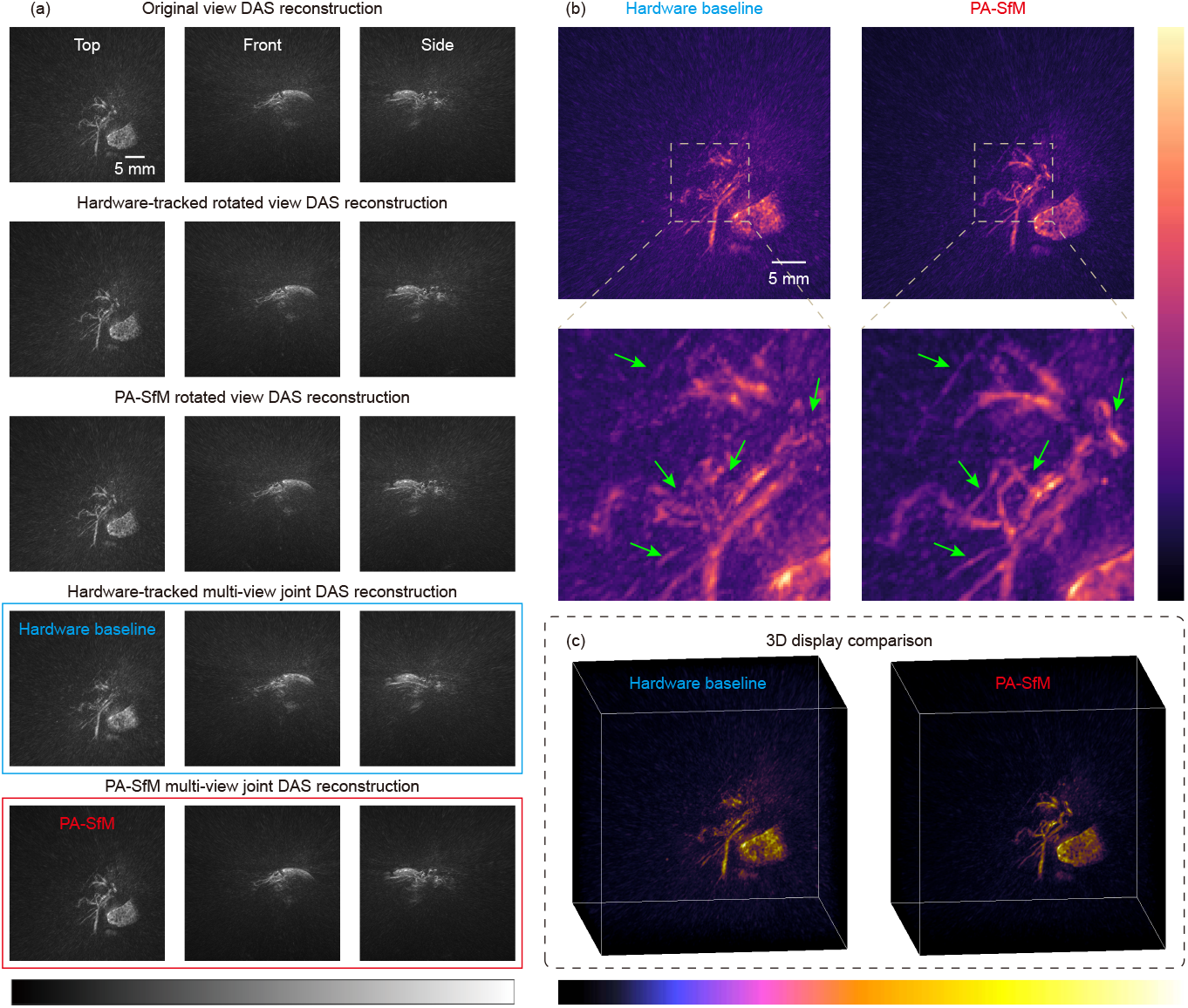
Multi-view PA reconstructions of mouse abdomen. (a) Top-, front-, and side-view MAPs of reconstructions from single views (original, hardware-tracked and PA-SfM rotated) and multi-views (hardware-tracked baseline, and PA-SfM result) (Scale: 5 mm; grayscale; reconstruction with DAS). (b) Locally magnified top-view MAPs comparing the hardware-tracked baseline and PA-SfM fine-tuned reconstruction (Scale: 5 mm; colormap: magma). (c) 3D display comparing hardware-tracked and PA-SfM reconstructions.

### 2.3 PA-SfM validation in animal study with translational multi-view reconstruction

In addition to rotational multi-view fusion, we further validated the performance of PA-SfM under translational multi-pose acquisition, which is a routine way to increase FOV. Using the same 3D PAI system, the *in vivo* imaging data is from a mechanical scanning experiment with four poses, as illustrated in Fig. 5(a). Starting from the initial position, the hemispherical array was mechanically translated three times, with a step size of 4 mm. Therefore, the four acquired poses jointly covered a larger anatomical region than a single stationary acquisition. For comparison, we reconstructed each individual pose using DAS and then performed multi-pose joint DAS reconstruction using either the hardware-tracked transformations or the transformations estimated by PA-SfM, as shown in Fig. 5(b), which also demonstrates that the single-pose reconstructions capture only partial vascular structures.

**Fig. 5.**
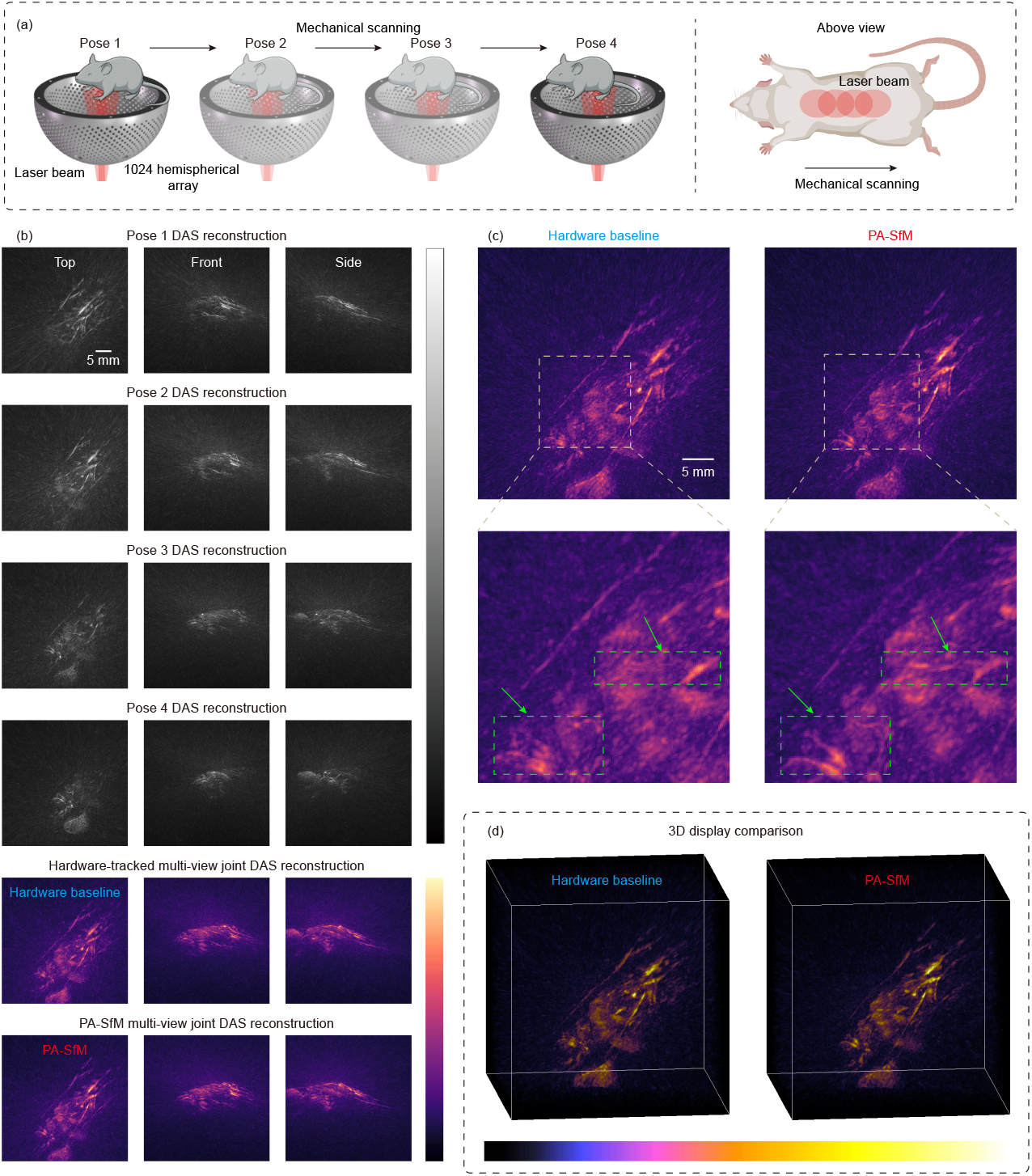
Expanded field-of-view multi-pose PA mapping using mechanical translation. (a) Schematic of the mechanical scanning experiment with the 1024-element hemispherical array. The animal was imaged at four poses, including the initial pose and three subsequent mechanical translations. Each translation step was 4 mm along a direction forming a 48° angle with the negative Y-axis of the array. (b) Top-, front-, and side-view MAPs of single-pose DAS reconstructions and multi-pose joint DAS reconstructions using hardware-tracked poses and PA-SfM estimated poses (Scale: 5 mm; colormap: gray, magma). (c) Top-view MAP comparison between the hardware-tracked baseline and PA-SfM reconstruction, with local magnifications highlighting vessel continuity and structural sharpness (Scale: 5 mm; colormap: magma). (d) 3D display comparison of the hardware-tracked and PA-SfM multi-pose reconstructions.

The advantage of PA-SfM is further highlighted in the comparison of top-view MAP results, as shown in Fig. 5(c). In the hardware-tracked baseline, several vessels appear fragmented or duplicated because of imperfect spatial alignment. The hardware-tracked multi-pose reconstruction expands the visible region by combining information from all poses; however, residual pose errors introduce noticeable blurring, vessel broadening, and discontinuities, especially for fine vascular structures. In contrast, the PA-SfM reconstruction substantially reduces these artifacts and restores clearer vessel structures, as indicated by the green arrows and dashed boxes. The 3D visualization in Fig. 5(d) also confirms that PA-SfM improves volumetric consistency after multi-pose joint reconstruction. These results demonstrate that PA-SfM enables multi-pose photoacoustic mapping without extra pose and position tracking techniques, thereby providing a powerful algorithm for tracker-free, freehand 3D PAI systems.

### 2.4 Quantitative validation in multi-view sparse-array simulations

Previous human and in vivo animal studies demonstrated the high performance of PA-SfM compared with mechanical position tracking. To further quantitatively evaluate pose recovery and image reconstruction under fully controlled conditions, we conducted numerical simulations using a phantom liver model under a multi-view setup. As shown in Fig. 6(a), the simulation utilized a sparse hemispherical array to acquire data from multiple views. Specifically, the array consisted of only 33 sparse elements distributed across a spherical cap with a radius of 60 mm. View 1 served as the reference array pose, while View 2 and View 3 were generated by translating and rotating the array around the target. The single view and PA-SfM multi-view joint reconstruction results are shown in Fig. 6(b,c).

**Fig. 6.**
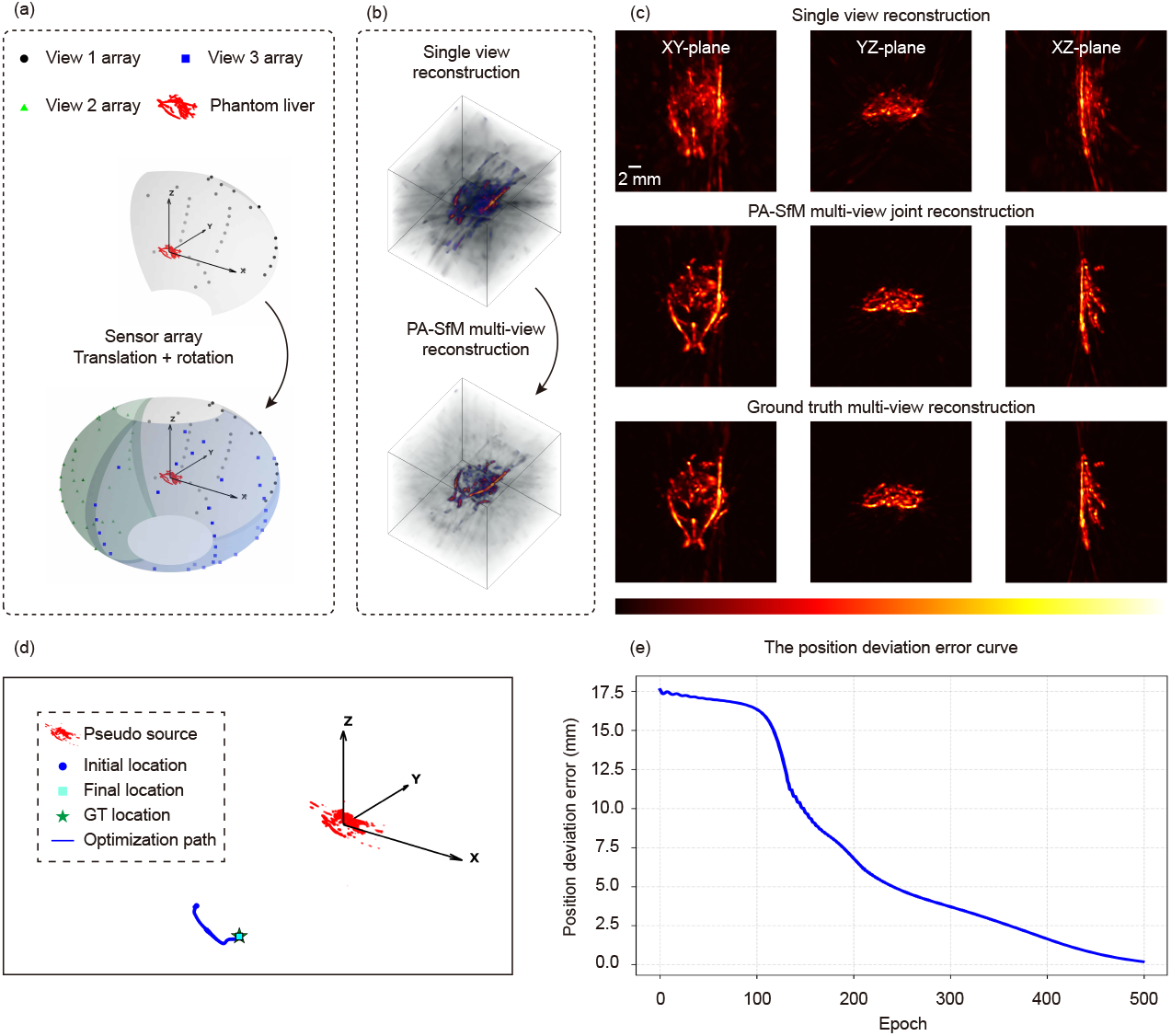
The reconstruction results of multi-view simulation data with PA-SfM algorithm and the PA signal of the target sensor at different stages. (a) The multi-view setup of the simulation experiment. (b) The 3D results of single-view, and PA-SfM multi-view reconstruction. (c) The single-view reconstruction, multi-view reconstruction using the PA-SfM pose, and GT of the phantom liver (Scale: 2 mm). (d) Differentiable single-sensor localization optimization. (e) The absolute position deviation error curve of the single-sensor localization optimization.

To demonstrate the pose optimization mechanism, Fig. 6(d-e) illustrates a representative target sensor during single-sensor localization process. Starting from an initial sensor location, the differentiable localization procedure progressively optimized the sensor position by minimizing the discrepancy between the simulated PA signal and the ground truth (GT) supervision signal. As evidenced by the absolute position deviation error curve between the predicted location and GT location of a randomly selected sensor in Fig. 6(e), the localization error decreased steadily during optimization, indicating effective convergence of the proposed differentiable sensor localization strategy. After PA-SfM pose optimization, the mean and minimum Euclidean distance errors of sensor localization were reduced to 0.751 mm and 0.113 mm, respectively, demonstrating accurate recovery of the multi-view array poses.

Based on the recovered PA-SfM poses, we further compared the 3D reconstruction results. As depicted in Fig. 6(b,c), the single-view reconstruction suffered from severe limited-view artifacts, yielding quantitative metrics of 25.51 dB for the peak signal-to-noise ratio (PSNR) and 0.6775 for the structural similarity index measure (SSIM) in the MAP along the XY plane. In contrast, the PA-SfM multi-view reconstruction effectively expanded the angular coverage and substantially suppressed limited-view artifacts. The reconstructed structures were highly consistent with the GT phantom liver, achieving a markedly improved PSNR of 41.14 dB and an SSIM of 0.9799. These results quantitatively validate that the proposed PA-SfM framework can accurately recover multi-view array poses and substantially improve 3D PA image reconstruction quality under sparse-array conditions.

### 2.5 In vivo animal study for validation of multi-view PA-SfM joint reconstruction with known relative geometry

Besides simulation quantitative assessment, we continued to validate PA-SfM on real *in vivo* animal imaging data with a known relative geometry between views. Specifically, we used data from *in vivo* multi-view 3D PAI of rat kidney (Fig. 7) and rat liver (Fig. 8), which were acquired by Professor Kim’s lab using a hemispherical ultrasound (US) transducer array with 1024 elements and a radius of 60 mm, and the average center frequency of each US transducer element in the array is 2.02 MHz [38]. More details regarding the 3D PAI system and animal experiments can be found in the literature [39–45].

**Fig. 7.**
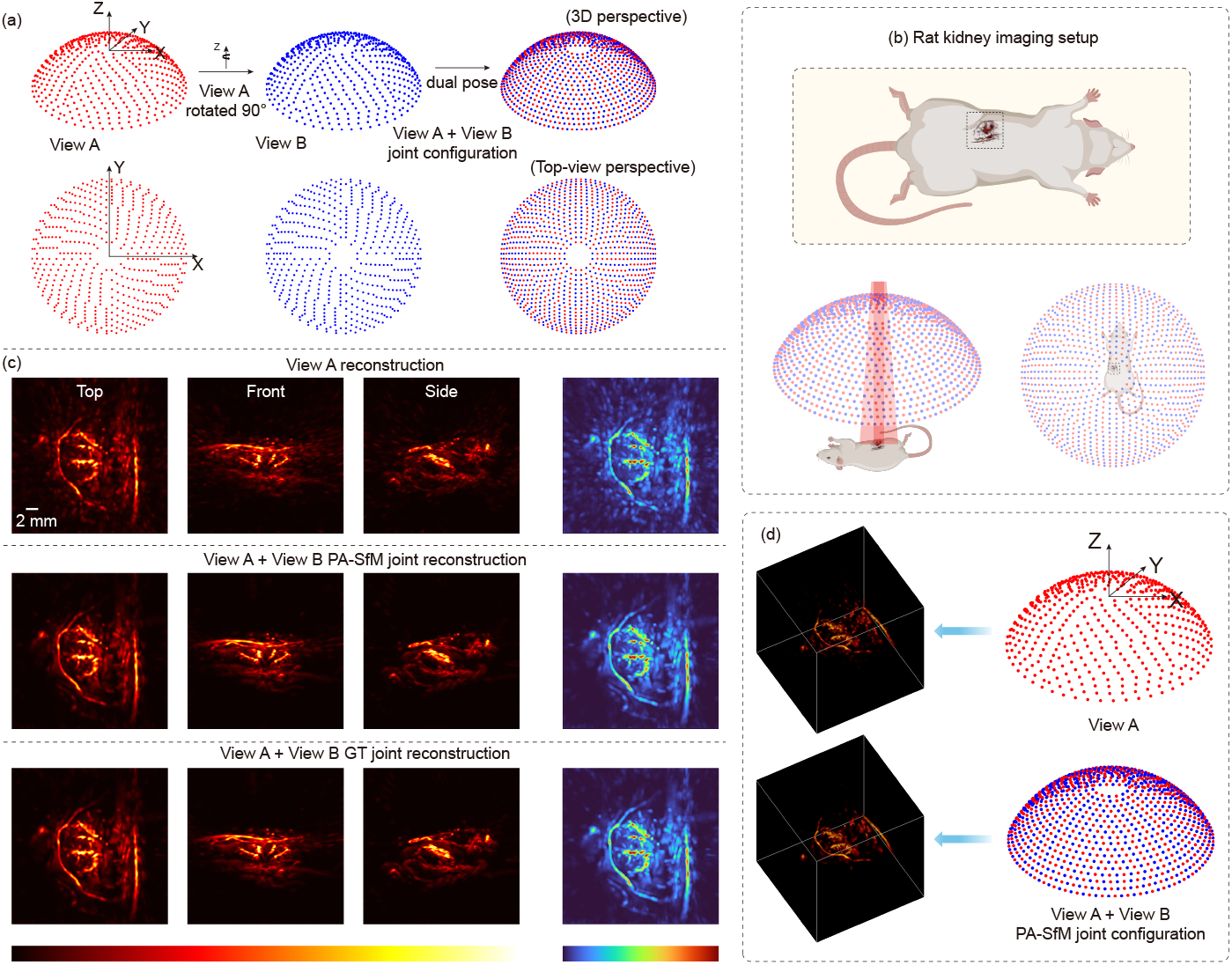
The reconstruction results of multi-view *in vivo* rat kidney with PA-SfM algorithm. (a) The dual-view 3D PAI for *in vivo* rat kidney imaging. (b) The rat kidney imaging setup. (c) The single-view reconstruction, multi-view reconstruction using the PA-SfM pose, and GT of the *in vivo* rat kidney (Scale: 2 mm). (d) The 3D results of single-view, and PA-SfM multi-view reconstruction.

**Fig. 8.**
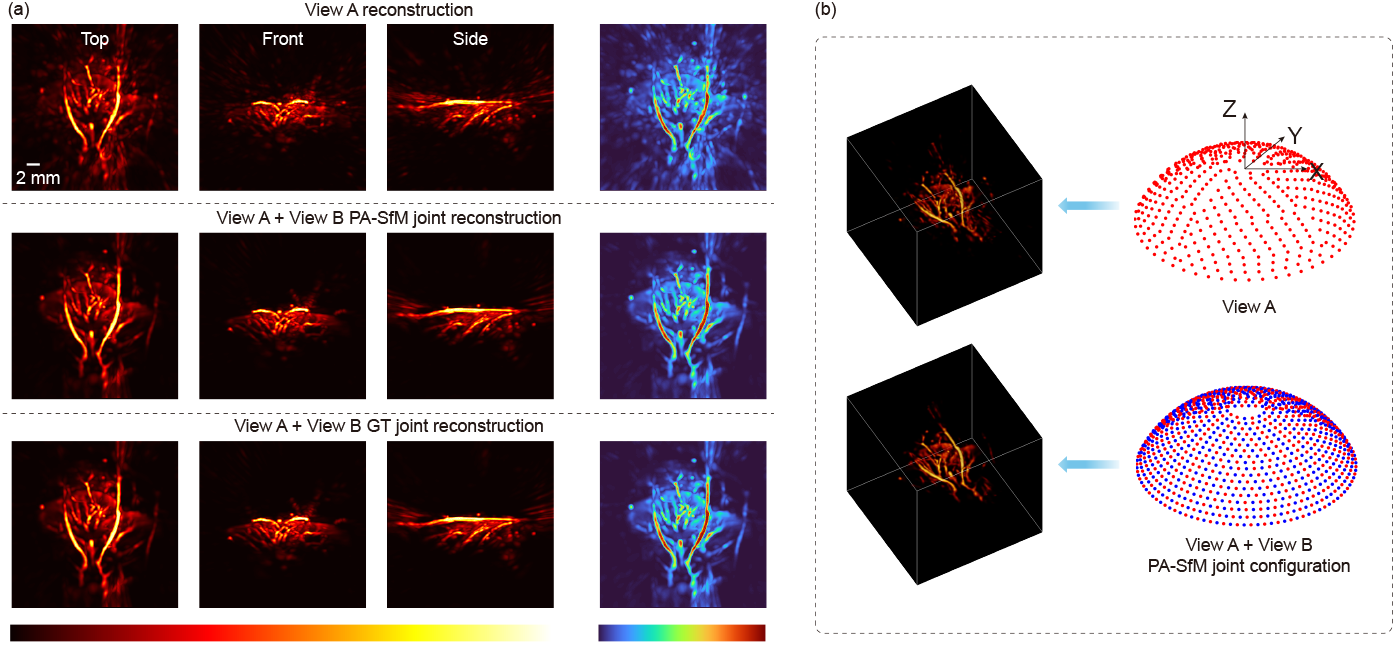
The reconstruction results of multi-view *in vivo* rat liver with PA-SfM algorithm. (a) The single-view reconstruction, multi-view reconstruction using the PA-SfM pose, and GT of the *in vivo* rat liver (Scale: 2 mm). (b) The 3D results of single-view, and PA-SfM multi-view reconstruction.

Based on the arrangement pattern of array elements, we deliberately partitioned the full 1024-element spherical array into two 512-element sparse subsets, separated by an exact 90° relative rotation along the azimuthal axis (Fig. 7(a-b)). These two subsets, designated as View A and View B which represent two independent imaging poses, were used to generate the GT multi-view reconstruction for quantitative evaluation. The final *in vivo* 3D reconstruction results are presented in Fig. 7(c-d).

For the rat kidney experiment, Fig. 7(c) compares the single-view reconstruction, the PA-SfM multi-view reconstruction, and the GT multi-view reconstruction. The MAPs from different viewpoints are shown to comprehensively evaluate the recovered 3D vascular structures, and the top-view MAP with a turbo colormap is further provided to highlight background noise and reconstruction artifacts. As shown in Fig. 7(c-d), the single-view reconstruction provides limited structural information and exhibits prominent limited-view artifacts. In contrast, the PA-SfM multi-view reconstruction successfully fuses the multi-view data from View A and View B, effectively overcoming the limited-view constraint and recovering a more complete 3D vascular network of the *in vivo* rat kidney. The reconstructed structures show high consistency with the GT multi-view reconstruction, demonstrating the capability of PA-SfM to recover accurate relative view geometry from real biological PA data.

Quantitative assessments based on the top-view MAP further corroborate these visual improvements. When evaluated against the GT multi-view reconstruction, the single-view kidney reconstruction yielded a PSNR of 30.20 dB and an SSIM of 0.7019. By contrast, the PA-SfM multi-view reconstruction achieved a substantially improved PSNR of 41.42 dB and an SSIM of 0.9864, indicating that the PA-SfM recovered pose enables accurate cross-view fusion and significantly suppresses motion-induced reconstruction artifacts. A further comparison between the PA-SfM recovered pose and the GT pose showed that the mean and minimum localization errors for all sensors were 0.34 mm and 0.027 mm, respectively, demonstrating the high accuracy of pose recovery.

To further demonstrate the robustness and generalizability of the proposed algorithm, we validated the method using the *in vivo* rat liver imaging data. Following the same visualization scheme, Fig. 8(a-b) presents the liver reconstruction results. The single-view liver reconstruction suffers from incomplete vessel connectivity and elevated background noise, which is particularly evident in the turbo colormap. In comparison, the PA-SfM multi-view reconstruction effectively integrates complementary information from the two views, restores continuous vascular structures, and recovers intricate vascular branching patterns that are highly consistent with the GT reconstruction.

These morphological observations are further supported by the quantitative metrics. Compared with the GT multi-view reconstruction, the single-view liver MAP yielded a PSNR of 28.01 dB and an SSIM of 0.6947. The PA-SfM multi-view reconstruction markedly improved the reconstruction accuracy, achieving a PSNR of 44.31 dB and an SSIM of 0.9961. Under the PA-SfM estimated array poses, the mean and minimum localization errors across all sensors were 0.21 mm and 0.016 mm, respectively. These *in vivo* results confirm that PA-SfM can accurately estimate multi-view relative poses and substantially improve 3D PA image reconstruction quality across different biological organs under sparse-view acquisition conditions.

## 3 Discussion

PA-SfM establishes a tracker-free route for freehand 3D PAI with superior high accurate pose recovery and joint reconstruction in multi-view, substantially reducing artifacts and expanding FOV. Rather than relying on motor encoders, scanning stages or external tracking devices, PA-SfM estimates the relative imaging geometry directly from the acquired PA measurements and incorporates the recovered poses into joint volumetric reconstruction. We demonstrated this principle in simulations, *in vivo* rat studies with known relative geometry, mechanically rotated and translated array acquisitions, and genuine freehand imaging of human hand vasculature. In the hand imaging experiment, PA-SfM has demonstrated its power to recover array pose and reconstruct a coherent large FOV from multi-view imaging with freehand motion without predefined motion trajectories. These results indicate that PA-SfM can turn otherwise uncontrolled motion into useful geometric diversity for 3D PAI.

Compared with mechanical tracking, PA-SfM improves reconstruction quality with more accurate pose estimation for the 3D PAI system used in this study. This improvement arises because the nominal encoder positions could be influenced by calibration errors, backlash, structural deviations or stochastic motion, leading to image blurring for fine structures. By optimizing the imaging geometry from PA signals themselves, PA-SfM effectively reduced these misregistration artifacts and produced sharper, more continuous fine vascular structures. The repeatability study in both human hand and animal studies further supported the reliability of the method. Nevertheless, the method still requires sufficient structural overlap and recognizable PA features across views, and its performance could degrade for sparse absorbers, weak contrast, severe artifacts or limited overlap. In addition, during expanded FOV freehand imaging, the array poses near the periphery of the overall FOV can receive relatively weaker optical illumination than those near the central region, resulting in reduced signal intensity close to edges in the joint reconstruction. We therefore applied edge compensation to the results shown in Fig. 1(e) and Fig. 2(c); further discussion is provided in Supplementary Note 1. The current formulation also assumes rigid relative geometry and does not explicitly model non-rigid tissue deformation, which should be addressed in future work for applications involving larger tissue motion or deformation.

Computational efficiency and reproducibility are also central to practical deployment of PA-SfM. We have therefore released the complete automated workflow for the continuous 10-frame PA-SfM pose estimation and joint reconstruction example in Fig. 2(c), together with the corresponding 10-frame dataset; detailed instructions are provided in Supplementary Note 2. In the current implementation of PA-SfM, pose recovery and joint reconstruction for each additional pose require only 175 s on a single NVIDIA A100 SXM4 40 GB GPU, and the complete 10-pose workflow takes less than 28 minutes, using a single reconstruction grid of 400 *×* 400 *×* 400 voxels. The GPU memory usage during the total process is approximately 5.1 GB. Although this implementation is currently suitable for offline processing, the runtime is relatively efficient for high-resolution volumetric 3D PAI, particularly because the framework jointly performs unknown pose estimation and volumetric reconstruction without external tracking, encoder feedback or predefined motion. To the best of our knowledge, PA-SfM is the first open-source computational framework in 3D PAI that integrates pose recovery and joint reconstruction into a unified pipeline. Its computational efficiency, modest GPU memory requirements, and cross-platform applicability facilitate adoption across a range of 3D PAI systems without additional tracking hardware. Future work will focus on improving computational speed through algorithmic acceleration, memory optimization and parallel implementation, with the goal of extending PA-SfM towards real-time pose recovery for interactive freehand 3D PAI.

## 4 Methods

The proposed PA-SfM framework is a closed-loop pipeline from reference reconstruction to unknown pose estimation, culminating in joint multi-view reconstruction (Fig. 9). The system is formulated upon a differentiable acoustic radiation engine and rigid-body geometry constraint. The detailed algorithmic workflow is structured into the following interacting modules.

**Fig. 9.**
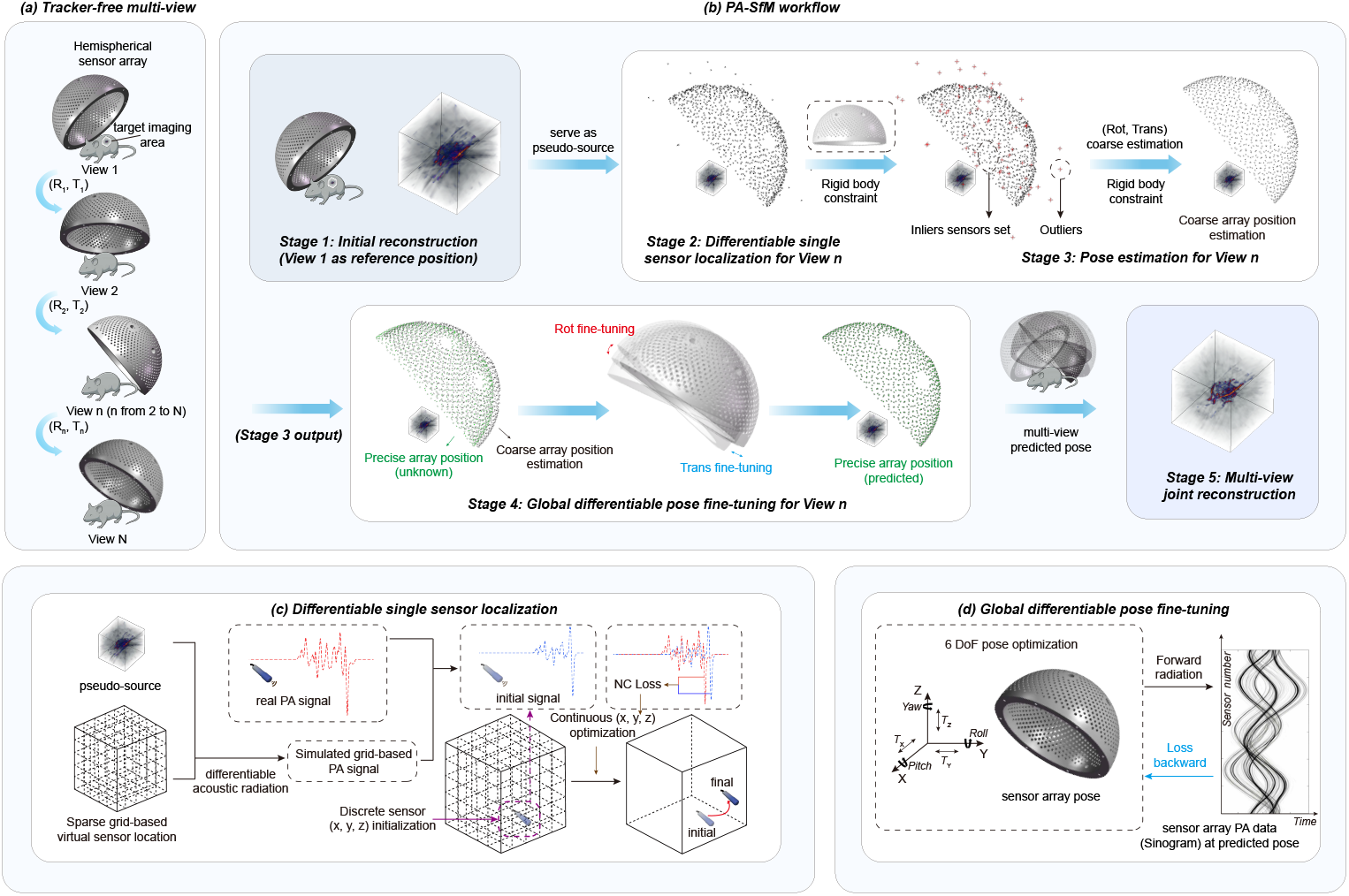
The overview framework of PA-SfM algorithm for tracker-free freehand 3D PAI. (a) The tracker-free multi-view imaging operation. (b) The PA-SfM pipeline. (c) The differentiable single probe localization of PA-SfM. (d) The global differentiable pose fine-tuning of PA-SfM.

### Differentiable Acoustic Radiation

The core of the framework is a differentiable acoustic radiation engine that models acoustic wave propagation and computes gradients. To map the 3D PA initial pressure distribution 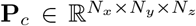to the time-resolved sensor signals *p*(**x**_*s*_, *t*) at sensor location **x**_*s*_ *∈* R^3^, we employ a Gaussian kernel-based propagation model:

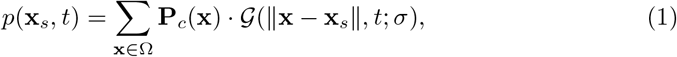

where Ω denotes the reconstructed volume, and *G* is the spatiotemporal Gaussian radiation kernel parameterized by width *σ*.

The Gaussian radiation kernel is derived from the analytical solution of the PA wave equation. The pressure field generated by an initial pressure distribution *p*_0_(**r**^*′*^) can be expressed using Green’s function as [46, 47]

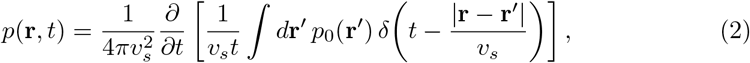

where **r** is the observation position, **r**^*′*^ is the source position, *v*_*s*_ is the speed of sound, and *p*_0_(**r**^*′*^) is the initial pressure distribution.

For a compact, spherically symmetric PA source element with radial initial pressure distribution *p*_0_(*r*), the analytical solution of the PA wave equation can be further written as

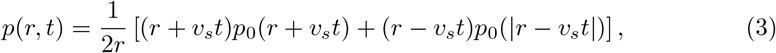

where *r* is the propagation distance (the detailed derivation is provided in Supplementary Note 3). Under the far-field condition, where the propagation distance is larger than the characteristic source size, the inward-propagating component becomes negligible and the detected signal is dominated by the outward-propagating component:

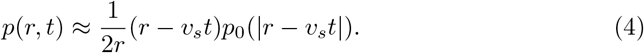

In this work, each voxel is modeled as a local Gaussian source element which has been demonstrated to be effective for 3D PA reconstruction [48], *p*_0_(*r*) = *p*_*c*_ exp (*r*^2^*/*2*σ*^2^) . Substituting this profile into the far-field expression gives

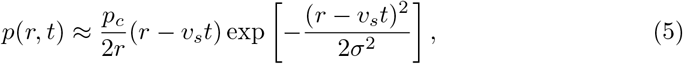

which motivates the Gaussian radiation kernel *G* used in Eq. (1). The detailed derivation from the photoacoustic wave equation to the far-field spherically symmetric solution, the Gaussian ball source approximation, and the GPU-accelerated far-field discrete model used in our engine is provided in Supplementary Note 3.

In the discrete implementation, each sensor’s temporal signal is the superposition of contributions from all independent Gaussian radiation sources according to its propagation distance and initial pressure value. This formulation exposes two natural parallel dimensions, sensor index and temporal index, and is therefore well suited for GPU execution. The forward and backward propagation operators are implemented as custom OpenAI Triton kernels and wrapped by `torch.autograd.Function`. As a result, gradients can be propagated not only to the source intensity **P**_*c*_ for image reconstruction, but also to the continuous sensor coordinates **x**_*s*_ for differentiable sensor localization and global pose refinement.

The main PA-SfM optimization workflow is summarized in Algorithm 1, with multi-view extension implemented by sequentially repeating the pose recovery and joint reconstruction steps for additional views.

### Stage 1: Initial Reference Reconstruction

The first stage is to establish a structural reference map (coarse model) using data acquired at a known, fixed pose (Pose 1 or View 1). Given the known sensor array coordinates **X**_1_ and the corresponding detected temporal signals **S**_1_, the initial source distribution **P**_*c*_ is initialized on a voxel grid. The reconstruction is formulated as an inverse optimization problem:

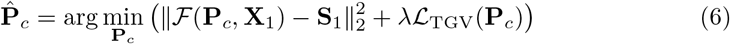

where *F* represents the forward simulation engine, and *L*_TGV_ is the Total Generalized Variation (TGV) regularization term utilized to preserve structural edges while suppressing background noise. The objective function is iteratively minimized using the Adam optimizer, yielding the initial 3D anatomical structure serving as the reference map.

### Stage 2: Differentiable Single Sensor Localization

To estimate the unknown sensor pose (Pose 2 or View 2), we independently localize each array element using the reference map 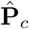 obtained in Stage 1. By fixing 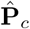, the individual sensor coordinates **x**_*s,i*_ become the optimizable parameters. The localization is driven by a Negative Correlation (NC) loss:

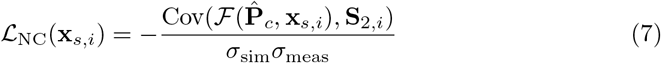

where *σ*_sim_ and *σ*_meas_ denote the standard deviations of the simulated and measured signals, respectively.

To circumvent local minima, a coarse-to-fine search strategy is employed, as demonstrated in Fig. 9(c):

1. **Global Coarse Search:** The target space is discretized into grids, and candidate locations are ranked based on *L*_NC_ to select the Top-*K* priors.
2. **Gradient-based Refinement:** Starting from the candidate priors, gradient descent is utilized to optimize the (*x, y, z*) coordinates.
3. **Dynamic Smoothing:** The Gaussian kernel width *σ* in Eq. (1) is dynamically decayed (from large to small) during optimization, acting as a simulated annealing mechanism to ensure convergence to the global optimum.

This stage outputs a set of independently predicted, albeit noisy, sensor coordinates 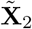

### Stage 3: Robust Rigid Array Estimation

The reference map **P**_*c*_ obtained in Stage 1 contains noise and many artifacts due to sparsity, leading to errors in sensor localization in Stage 2. To suppress these errors, the inherent geometric constraints of the rigid sensor array are leveraged. Given the predefined geometric template of the array 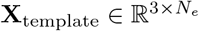 (where *N*_*e*_ is the number of sensor elements), the task is transformed into finding the optimal global rotation matrix **R** *∈ SO*(3) and translation vector **t** *∈ ℝ*^3^.

A modified Random Sample Consensus (RANSAC) [49] algorithm is implemented.

To drastically improve search efficiency, a geometric consistency pre-check is introduced: randomly sampled point sets are evaluated based on their pairwise edge distances before performing Singular Value Decomposition (SVD). For valid subsets, the Kabsch algorithm [50] is applied to solve the rigid transformation. This stage yields a corrected global array coordinate set 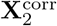 and reliably identifies a subset of inlier sensors *I*, whose relative positions satisfy the rigid array constraint within a preset small error tolerance.

### Stage 4: Global Differentiable Fine-tuning

To further enhance spatial precision, a RigidArrayOptimizer module is constructed. The optimization variables are globally parameterized as Euler angles ***θ*** and the translation vector **t** (Fig. 9(d)), ensuring that the array strictly adheres to rigid body kinematics: **X**_2_(***θ*, t**) = **R**(***θ***)**X**_template_ + **t**.

We introduce an inlier-masked gradient flow mechanism. The global loss is computed exclusively on the highly reliable signals from the inlier set *I* (identified in Stage 3):

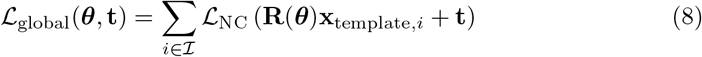

Crucially, while the loss is masked, the gradients backpropagate to update the global parameters (***θ*, t**). This mechanism physically compels the unreliable (outlier) sensors to be guided into their correct anatomical positions by the reliable (inlier) sensors, yielding highly precise Pose 2 coordinates. The importance of this stage is substantiated through quantitative experimental evaluation in section 2.4 and section 2.5. The detailed analysis of the intermediate coarse pose estimation and global differentiable pose fine-tuning is provided in Supplementary Note 4 and Supplementary Note 5.

### Stage 5: Joint Dual-view Reconstruction

In the final stage, the data from the known Pose 1 (**X**_1_, **S**_1_) and the calibrated Pose 2 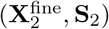 are seamlessly fused to form an augmented synthetic aperture. The fused dual-view data can be jointly reconstructed using either back-projection reconstruction or model-based iterative reconstruction methods. Therefore, the sparsity artifacts in the single view are effectively mitigated by coherently combining the dual-view data.

### Stage 6: Joint Multi-view Reconstruction

For scenarios involving more than two imaging views, the framework adopts a sequential pose-chain calibration strategy to recover all sensor poses within a unified global coordinate system. The coordinate system of Pose 1 is defined as the global reference frame. After Pose 2 is localized and refined through Stages 2–4, its estimated rigid transformation is used to map the Pose 2 sensor array into the Pose 1 coordinate frame.

For each subsequent unknown view, the pose is estimated relative to the previously calibrated view rather than independently in isolation. Specifically, the relative rigid transformation between Pose *k* and Pose *k−* 1 is first recovered using the same differentiable localization, robust rigid estimation, and global fine-tuning pipeline described in Stages 2–4. The resulting relative transformation is then compounded with the accumulated transformations from previous stages, thereby expressing Pose *k* in the original Pose 1 coordinate system:

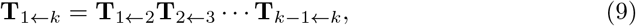

where **T**_*a←b*_ *∈ SE*(3) denotes the rigid transformation that maps coordinates from the Pose *b* frame to the Pose *a* frame.

By recursively chaining these relative transformations, all sensor arrays from multiple views are consistently registered into the initial Pose 1 coordinate frame. The calibrated coordinates from all poses are then combined to form a large synthetic sensing aperture:

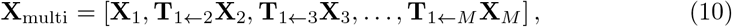

where *M* is the total number of imaging views. Finally, a comprehensive joint reconstruction is performed using all calibrated sensor coordinates and their corresponding measurements. This unified multi-view reconstruction maximally expands the effective aperture, suppresses sparsity-induced artifacts, and yields the final high-fidelity 3D photoacoustic volume.

## Acknowledgements

This research was supported by the following grants: the National Key R&D Program of China (No. 2023YFC2411700, No. 2017YFE0104200); the Beijing Natural Science Foundation (No. 7232177); the National Natural Science Foundation of China (62472213); the Basic Science Research Program through the National Research Foundation of Korea (NRF) funded by the Ministry of Education (2020R1A6A1A03047902).

We would like to express our gratitude to Zhibo Xiao, Tingting Huang and Boxin Yao for their generous assistance.

## Algorithm 1

Differentiable Photoacoustic Structure from Motion (PA-SfM)

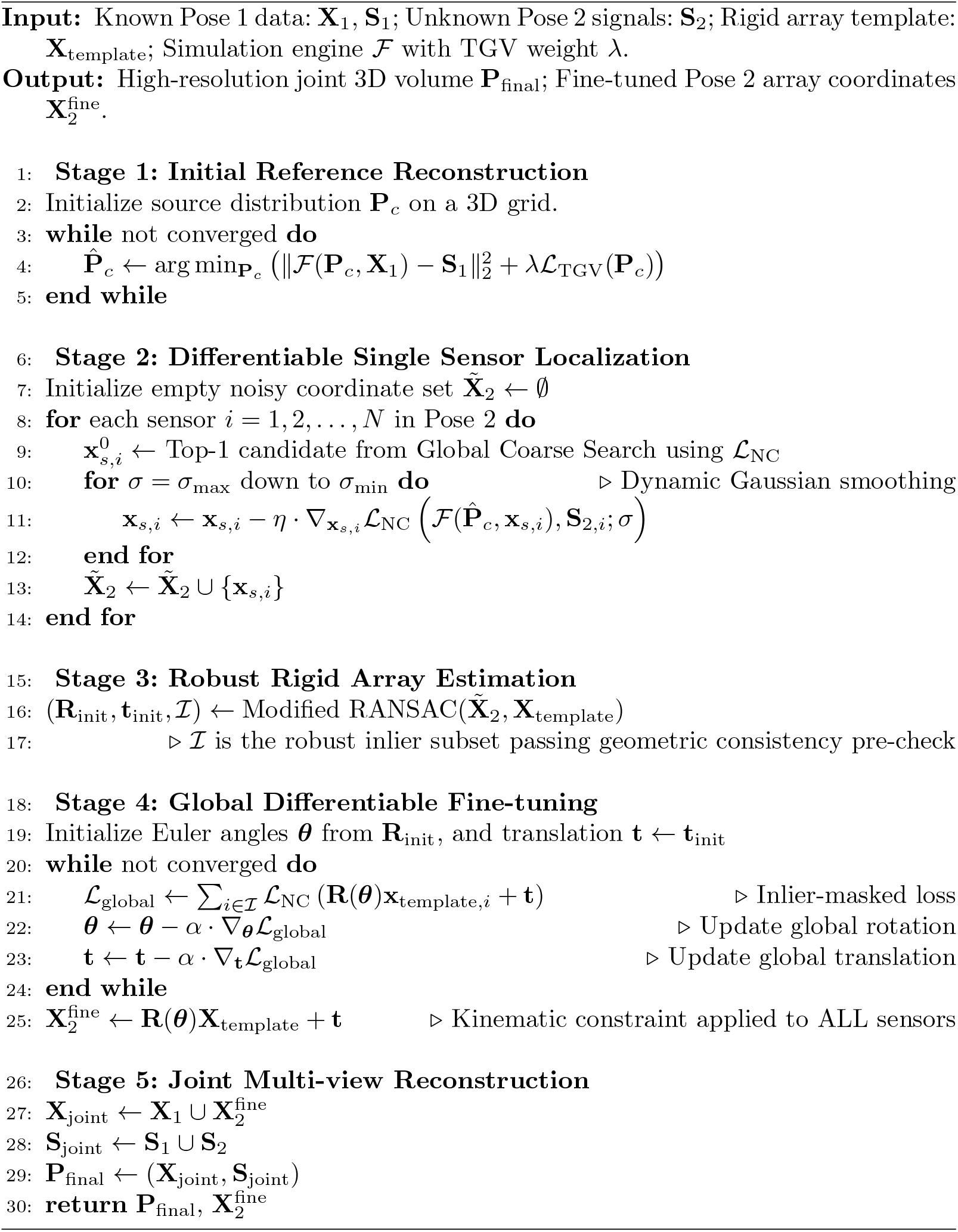

## Declarations

### Data availability

The data supporting the findings of this study are provided within the Article and its Supplementary Information. The datasets generated during the study are available for research purposes from the corresponding authors on reasonable request.

### Code availability

The source code for PA-SfM has been made publicly available in the following GitHub repository: https://github.com/JaegerCQ/PA-SfM.

### Author contributions

S.L. and J.G. conceived and designed the study and the algorithms. C.K., S.C., H.H., X.W. and J.S. provided the *in vivo* data. Q.C., Y.W., S.W. and Y.Z. contributed to the simulation experiments. Y.Z. contributed to the metrics calculation. C.L. and Y.Y. supervised the study. All of the authors contributed to writing the paper.

